# Hindlimb Unloading Induces Bone Microarchitectural and Transcriptomic Changes in Murine Long Bones in an Age-Dependent Manner

**DOI:** 10.1101/2023.10.09.561510

**Authors:** Steven J. Meas, Gabriella M. Daire, Michael A. Friedman, Rachel DeNapoli, Preetam Ghosh, Joshua N. Farr, Henry J. Donahue

**Author notes:** Corresponding Author: Henry J. Donahue. Contributions: Conceptualization: SJM, MAF, JNF, HJD Methodology: SJM, MAF, PG, JNF, HJD Investigation: SJM, GMD, RD Analysis: SJM, GMD, MAF, RD, PG, HJD Writing and Editing: SJM, GMD, MAF, RD, PG, JNF, HJD.

## Abstract

Age and disuse-related bone loss both result in decreases in bone mineral density, cortical thickness, and trabecular thickness and connectivity. Disuse induces physiological changes in bone like those seen with aging.

Bone microarchitecture and biomechanical properties were compared between 6- and 22-month-old C57BL/6J male control mice and 6-month-old mice that were hindlimb unloaded (HLU) for 3 weeks. Epiphyseal trabecular bone was the compartment most affected by HLU and demonstrated an intermediate bone phenotype between age-matched controls and aged controls.

RNA extracted from whole-bone marrow-flushed tibiae was sequenced and analyzed. Differential gene expression analysis additionally included 4-month-old male mice unloaded for 3 weeks compared to age-matched controls. Gene ontology analysis demonstrated that there were age-dependent differences in differentially expressed genes. Genes related to downregulation of cellular processes were most affected in 4-month-old mice after disuse whereas those related to mitochondrial function were most affected in 6- month-old mice. Cell-cycle transition was downregulated with aging.

A publicly available dataset (GSE169292) from 3-month female C57BL/6N mice unloaded for 7 days was included in ingenuity pathway analysis with the other datasets. IPA was used to identify the leading canonical pathways and upstream regulators in each HLU age group. IPA identified “Senescence Pathway” as the second leading canonical pathway enriched in mice exposed to HLU. HLU induced activation of the senescence pathway in 3- month and 4-month-old mice but inhibited it in 6-month-old mice.

In conclusion, we demonstrate that hindlimb unloading and aging initiate similar changes in bone microarchitecture and gene expression. However, aging is responsible for more significant transcriptome and tissue-level changes compared to hindlimb unloading.

**Highlights:**

- Epiphyseal trabecular bone is most susceptible to hindlimb unloading.
- Hindlimb unloaded limbs resemble an intermediate phenotype between age-matched and aged controls.
- Hindlimb unloading induces gene expression changes that are age dependent and may lead to inflammation and/or mitochondrial dysfunction depending on context.
- Younger mice (3-4 months) activate the senescence pathway upon hindlimb unloading, whereas skeletally mature (6 months) mice inhibit it.

## INTRODUCTION

Osteoporosis is a disease that is caused by low bone mineral density (BMD) and leads to an increased risk of fracture, with significant effects on morbidity and mortality. Osteoporosis increases in prevalence with age; hence the burden of osteoporotic fractures is predicted to increase as the global population ages[1]. Decreased mechanical loading, as observed in patients with limited mobility, e.g patients with hip fractures, can result in disuse-related bone loss[2]. An emerging concept is that disuse induces physiological changes in bone similar to those seen with aging[2,3]. Age and disuse-related bone loss both result in decreases in BMD, cortical thickness, and trabecular thickness and connectivity[2]. This effect on bone reduces biomechanical strength and fracture resistance[4]. Whereas age-related bone loss progresses at a rate of about 0.17% per month, disuse-related bone loss demonstrates an accelerated rate of bone loss at 1-2% per month[3,4]. Several strategies (i.e. weight-bearing exercise and resistance training) have been employed to combat both age- and disuse-related bone loss, with limited effectiveness[5]. Hence, new strategies for preventing the progression of both types of bone loss are needed to promote healthy aging.

One of the hallmarks of age-related bone loss is cellular senescence; a process characterized by cell cycle arrest and distinct changes in cellular expression, metabolism, and secretion[6]. Senescent cells accumulate in aging tissues and establish a senescence associated secretory phenotype (SASP), which is associated with detrimental effects on tissue[6]. In bone, SASP is associated with an increase in osteoclastogenesis and bone resorption, especially evident following a marked increase in number of senescent osteocytes in mice older than 18 months[7-10]. These age-related changes are paralleled in models of disuse, where unloading causes increased osteoclastogenesis and activity, decreased osteoblastogenesis, and increased bone marrow adiposity[2]. Despite recent advances in the field of bone biology and senescence, there has been little consideration of senescence in the context of disuse-induced bone loss, despite parallels to age-related bone loss, including increased inflammation[11-13]. The few studies that have considered senescence in the context of disuse examined the vertebral column or musculature surrounding the long bones[14,15]. However, the long bones themselves, primarily the femur and tibia, are significantly affected by disuse-paradigms such as hindlimb unloading or single limb immobilization[16,17].

To date, the relationship between senescence, age- and disuse-related bone loss has not been directly examined, especially in terms of the transcriptome. This is a significant gap in understanding the mechanisms disrupting bone homeostasis and the onset of unloading-induced osteoporosis. We hypothesize that age- and disuse-related bone loss develop through similar mechanisms, including senescence. Herein, we aimed to examine the novel role of senescence in disuse-related bone loss, and to directly compare the phenotype in age- and disuse-related bone loss.

## MATERIALS AND METHODS

### Animal Model

All animal procedures were approved by the VCU IACUC. Animal studies used 6- month and 22-month C57BL/6J male mice (000664, Jackson laboratories). Animals were dual-housed. The disuse-related model of bone loss was established by hindlimb unloading (HLU) for 3 weeks, as has been performed extensively in our laboratory [17-19]. These experiments were performed using rat cages with a wire mesh bottom (Allentown) to allow for tail suspension. Under anesthesia, mice tails were wrapped with two strands of porous medical grade tape (3M). The distal portion of the tape was attached to a steel eye hook, which was connected to string wrapped around an elevated crossbar. The line was rotated, elevating the tail and hindlimbs of the mouse until the hindlimbs formed an angle of 30º with the ground. This set-up provided mice free access to food and water within a restricted navigable area. If mice lost >10% body weight, they were supplemented with DietGel Boost (ClearH2O). Mice were euthanized if humane endpoints were reached (loss in body weight >20%, open-ulcers), and those mice were excluded from the study. Power calculation predicted a sample size of 5 mice was needed per group. Total group numbers for mice included 6-month-old male control (n=8), 6-month-old male HLU (n=8), and 22-month-old male control (n=11).

### microCT parameters

Before and after HLU, mice femurs were scanned using X-ray micro computed tomography (SkyScan, Bruker). Animals were anesthetized using vaporized isofluorane and subjected to a scan of the femur at a resolution of 4032 × 3024 and 7μm. Images were reconstructed in three dimensions and analyzed using software from the manufacturer (Bruker). A 180μm section of the midpoint was isolated for cortical bone analysis. A 720μm section was isolated proximal to the epiphyseal plate for metaphyseal trabecular bone analysis. A 520μm section was isolated distal to the epiphyseal plate for epiphyseal trabecular bone analysis. The following cortical bone parameters were examined: bone mineral density, cortical area fraction, and cross-sectional thickness. The following trabecular bone parameters were examined: percent bone volume, trabecular thickness, trabecular number, and trabecular separation. Bone parameters were analyzed using final values and as a percent change from baseline. Femurs were rescanned post-sacrifice if images could not be used for analysis, however samples were excluded from trabecular bone analysis if baseline images were not clear. Bone mineral density was calibrated based on attenuation coefficients from samples with known density. Attenuation coefficient for 0.25 g/cm^3^ was 0.019 cm^2^/g, and 0.75 g/cm^3^ was 0.036 cm^2^/g. Samples for any of these parameters were excluded from analysis if computational results were beyond mean ± 2 standard deviations or if they contradicted empirical evidence.

### Biomechanical testing

Mechanical properties of femurs were evaluated by three-point bending to failure under displacement control at 1.0mm/min using Bose ElectroForce 3200, as previously described[16]. The following parameters were examined: yield load, ultimate load, yield displacement, work-to-fracture, yield stress, ultimate stress, post-yield strain and total toughness.

### Transcriptomics

Bone marrow flushed tibias were stored in RNAlater (Sigma-Aldrich, R0901-500ML) overnight at 4ºC before removing solution and storing at -80ºC until ready for processing.

Frozen bones were ground with a mortar and pestle in liquid nitrogen and bone dust was homogenized in bead mill tubes (FisherBrand, 15-340-151) in Trizol (Thermo Fisher, 15596026). RNA was extracted from bone-marrow flushed tibias from each group using RNEasy Mini Kit (Qiagen, 74104) and samples with an RNA integrity number greater than 4.0 were used for bulk RNAseq. RNAseq library was prepared (Illumina stranded mRNA prep) and performed on Nextseq 2000 P3 – 200 cycles (up to 1.1 billion reads) (Illumina). Paired-end raw sequence data was processed using BioJupies and analyzed using plug-ins for analysis[20]. This online tool uses the kallisto pseudoaligner to generate gene counts. Primary component analysis, clustergrammer[21] were performed on the top 2500 genes, using logCPM normalized values. Differential gene expression was powered using limma [22].

We also performed analysis of RNASeq data using Ingenuity Pathway Analysis (IPA) from Qiagen to determine whether “Senescence Pathway” is an upregulated Canonical pathway in HLU mice[23]. We included two additional datasets in our comparison analysis. First, 4-month-old C57BL/6J male mice subject to HLU for a period of three-weeks from our recently published study, which included detailed analysis of bone geometry and mechanical properties[19]. Second, a dataset (GSE169292) from the NIH GEO which contained results from 3-month-old female C57BL/6N mice subjected to HLU for a period of seven days[24]. A summary of experimental groups can be found in Table 1. IPA was performed using approximately the top 3000 molecules with lowest p values from differential gene expression analysis of control versus HLU in each age group. P values were calculated by the software based on logFC values using the Fisher exact test to determine if the “Senescence Pathway” was enriched. Benjamini-Hochberg procedure to correct for the false discovery rate was calculated by the software and used to identify key upstream regulators.

**Table 1.**
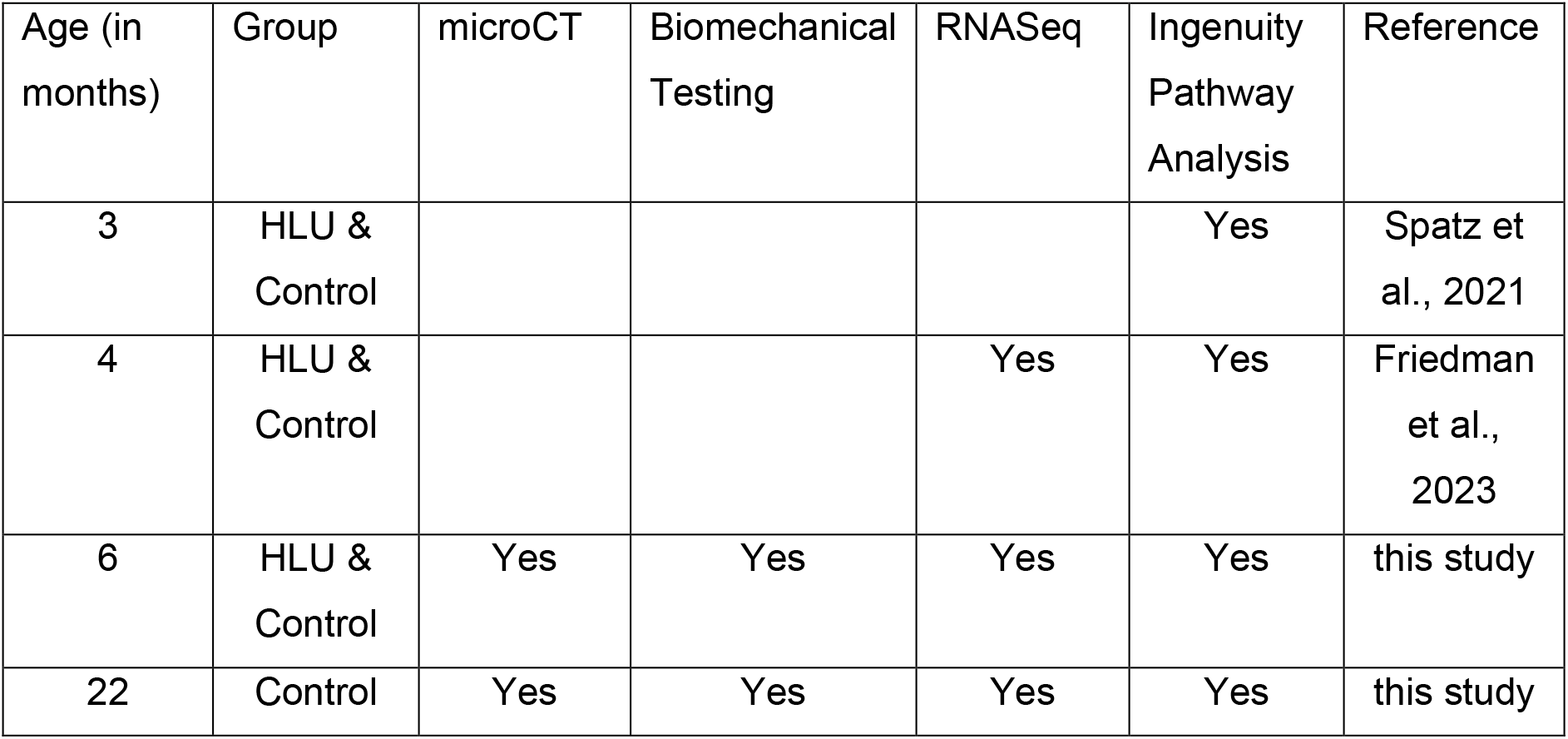
Summary of experimental groups.

| Age (in months) | Group | microCT | Biomechanical Testing | RNASeq | Ingenuity Pathway Analysis | Reference |
| --- | --- | --- | --- | --- | --- | --- |
| 3 | HLU & Control |  |  |  | Yes | Spatz et al., 2021 |
| 4 | HLU & Control |  |  | Yes | Yes | Friedman et al., 2023 |
| 6 | HLU & Control | Yes | Yes | Yes | Yes | this study |
| 22 | Control | Yes | Yes | Yes | Yes | this study |

### Statistical analyses

All statistical analyses for indices of bone microarchitecture and biomechanical testing were performed using GraphPad Prism (Version 9.5.1). Data was performed using a one-way ANOVA, α = 0.05, followed by Tukey’s post-hoc to adjust for multiple comparisons.

### Data Availability

RNAseq raw and processed files have been uploaded to GEO and are available under the following accession number: GSE235942.

## RESULTS

### Hindlimb unloading in a young skeletally mature mouse induces an intermediate phenotype between age-matched control mice and aged mice

After three weeks of the intervention period, there were no significant differences in cortical bone parameters in young control (n=8) and young HLU (n=8) mice (Figure 1A-D, S1).

**Figure 1.**
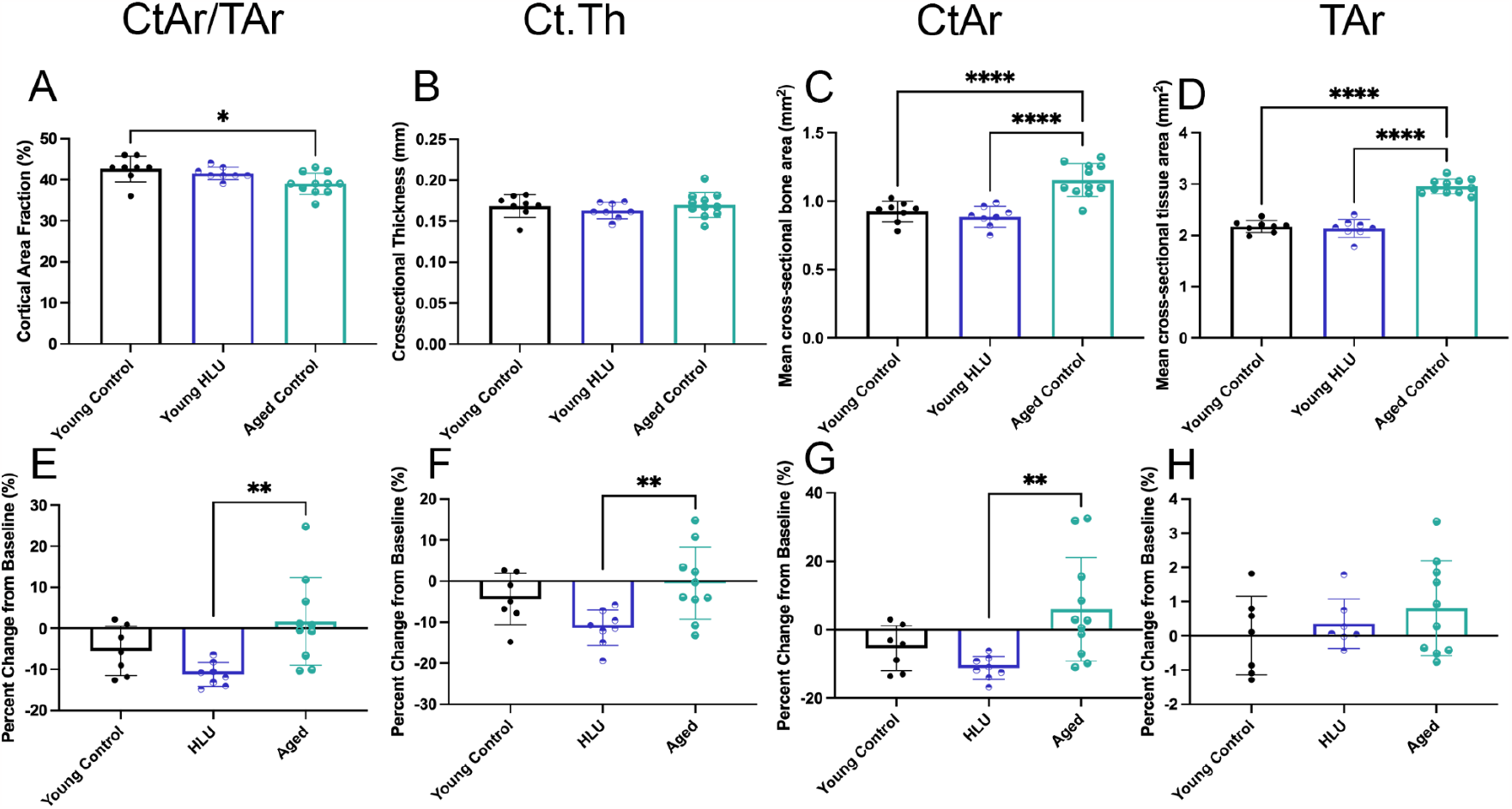
Differences in cortical bone parameters in male mice after the three-week intervention period. Top: absolute values. Bottom: percent change from baseline Cortical area fraction (CtAr/TAr: A,E), cortical thickness (Ct.Th: B, F), cortical bone area (CtAr: C, G), tissue area (TAr: D, H). HLU = hind-limb unloading * p < 0.05, ** p < 0.01, *** p < 0.01, **** p < 0.001, n = 8-11.

However, both cortical area (CtAr) and tissue area (TAr) were significantly increased when comparing the aged group (n=11) to either young group. Interestingly, cortical area fraction (CTAr/TAr) was significantly decreased when comparing the aged group (39%) to young control (42.63%, p < 0.05), but not to young HLU (41.50%). There were no significant differences in cortical thickness (Ct.Th) between groups. To determine the effects of the three-week intervention period we also calculated percentage change from baseline to characterize indices of bone loss (Figure 1E-F). From this perspective, both young control and HLU mice demonstrated a decrease in the four cortical bone indices, except for TAr, whereas aged mice demonstrated either little change or a slight increase. The HLU group was significantly different from the aged group when considering CtAr/TAr (-11.22% vs 1.691%, p < 0.01), Ct.Th (-11.39% vs -0.5516%, p < 0.01), and CtAr (-11.19% vs 5.964%, p < 0.01). There were no significant changes in TMD in male mice (Figure S2).

Similarly, to cortical bone, indices of metaphyseal trabecular bone were not significantly different between young control and young HLU groups (Figure 2A-D). Both young groups had lower trabecular thickness (Tb.Th) and trabecular separation (Tb.S) compared to aged controls. However, there were no significant changes in trabecular volume (BV/TV) and trabecular number (Tb.N). In terms of change from baseline (Figure 2E-F), BV/TV, Tb.Th and Tb.N declined in both control and HLU mice, and was significant when compared to aged mice which did not decrease as much. Tb.S increased in both control and HLU mice, but only HLU was significantly different from aged mice (+29.33% vs -4.113%, p < 0.01).

**Figure 2.**
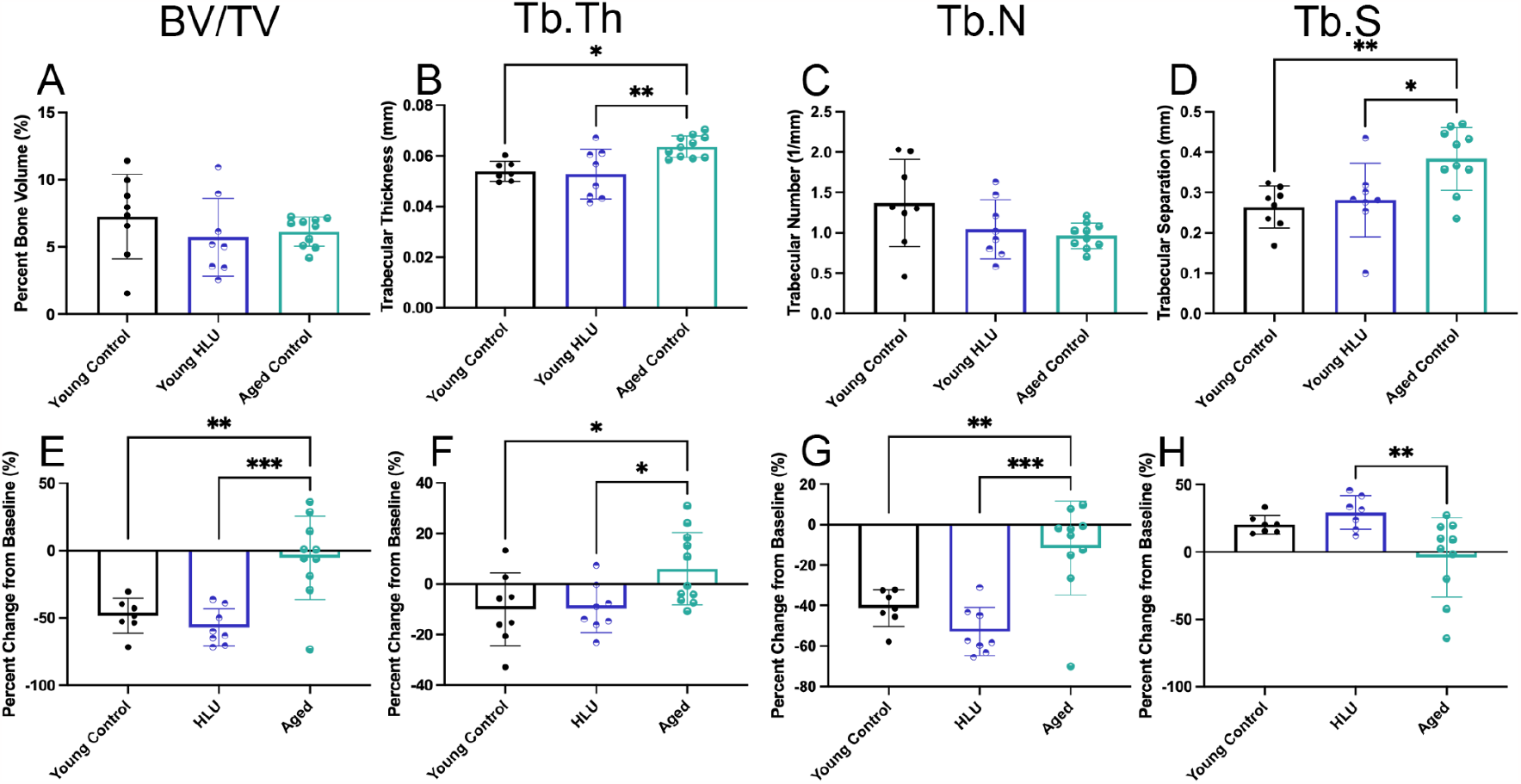
Differences in metaphyseal trabecular bone parameters in male mice after the three-week intervention period. Top: absolute values. Bottom: percent change from baseline. Trabecular bone volume (BV/TV: A,E), trabecular thickness (Tb.Th: B, F), trabecular number (Tb.N: C, G), trabecular separation (Tb.S: D, H). HLU = hind-limb unloading * p < 0.05, ** p < 0.01, *** p < 0.01, **** p < 0.001, n = 8-11

Epiphyseal trabecular bone appeared to be more susceptible to disuse than metaphyseal trabecular bone (Figure 3A-D). There was a decrease in BV/TV and Tb.Th in the HLU group compared to the young control group (BV/TV: 26.94% vs 39.93%, p < 0.05, Tb.Th: 0.06362 mm vs 0.07564 mm, p < 0.05). There was a decrease in BV/TV and Tb.N, and an increase in Tb.S in the aged group compared to the young control group (BV/TV: 27.46% vs 37.93%, p < 0.05, Tb.N: 4.038 1/mm vs 4.999 1/mm, p < 0.05, Tb.S: 0.1716 mm vs 0.1462 mm, p < 0.05). There were no significant differences between young HLU and aged groups. Considering change from baseline (Figure 3E-F), the young HLU demonstrated the greatest decline in BV/TV, Tb.Th and Tb.N, and increase in Tb.S. The young HLU group was significantly different compared to both young control and aged groups in all four epiphyseal trabecular bone parameters. This indicates that epiphyseal bone was the bone compartment most affected by disuse and was sufficient to accelerate the bone loss of a 6-month-old mouse to the degree of bone content to that of a 22-month-old mouse.

**Figure 3.**
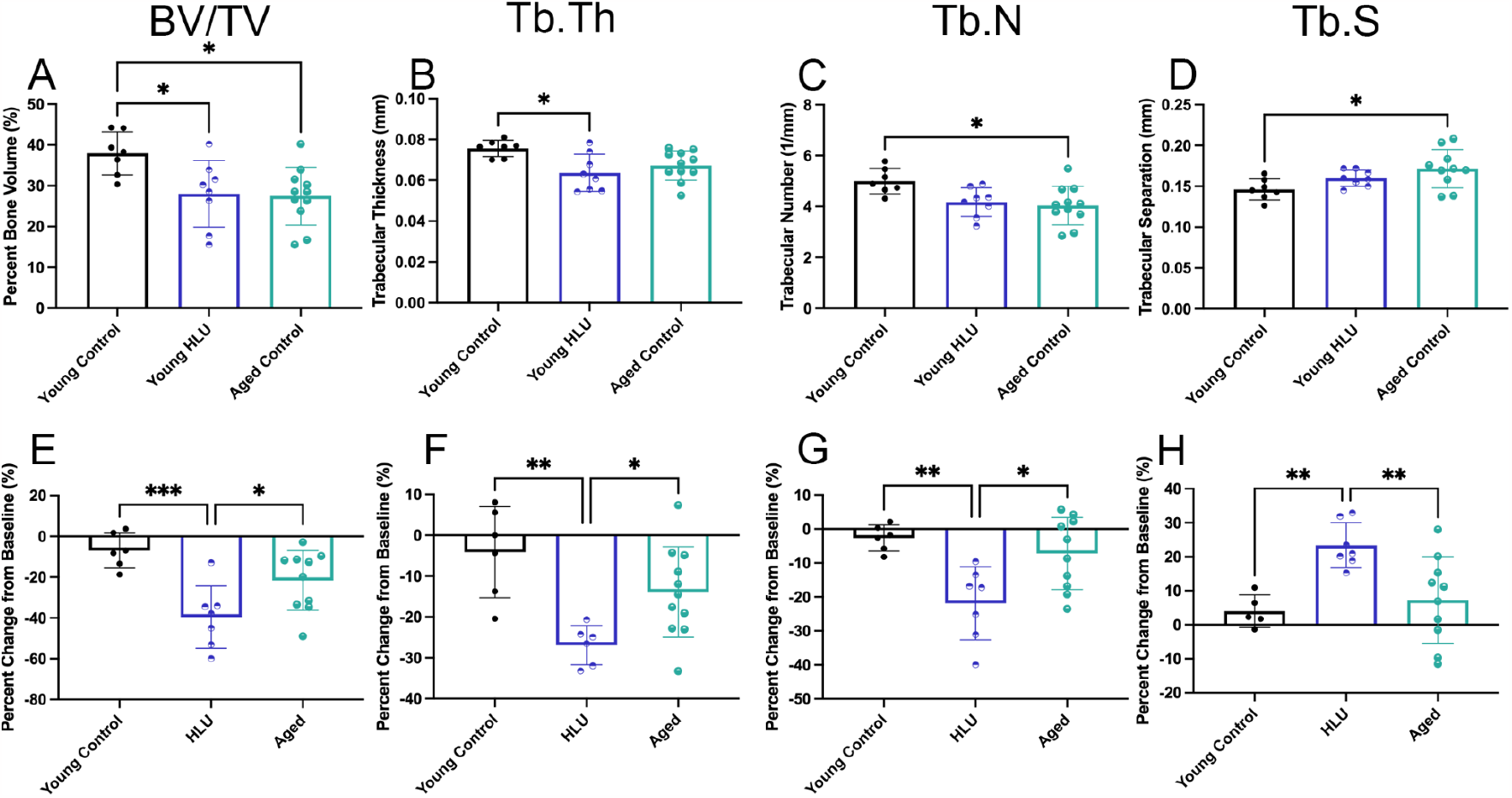
Differences in epiphyseal trabecular bone parameters in male mice after the three- week intervention period. Top: absolute values. Bottom: percent change from baseline. Trabecular bone volume (BV/TV: A,E), trabecular thickness (Tb.Th: B, F), trabecular number (Tb.N: C, G), trabecular separation (Tb.S: D, H). HLU = hind-limb unloading * p < 0.05, ** p < 0.01, *** p < 0.01, **** p < 0.001, n = 8-11.

### Femoral mechanical strength was different based on age in male mice

Three-point bending was performed on right femurs to compare bone strength between disuse and aging (Figure 4). Aged femurs demonstrated increased yield load and ultimate load, and decreased work-to-fracture compared to either young control or young HLU. Aged femurs also demonstrated a decrease in total toughness compared to young control, but young HLU demonstrated an intermediate phenotype. There were no significant differences in post-yield deformation, yield stress, ultimate stress or post-yield strain. These findings are consistent with microarchitecture results given that the primary correlates of bone strength in three-point bending are cortical area and cortical thickness of the mid- diaphysis[25]. These findings indicate that bones in male mice are weaker with age. Disuse initiates changes in bone strength, but perhaps three weeks are not sufficient to observe large changes in mechanical strength at these age groups. In 4-month-old mice there are changes in ultimate force after HLU, but no significant changes in any other parameter[19].

**Figure 4.**
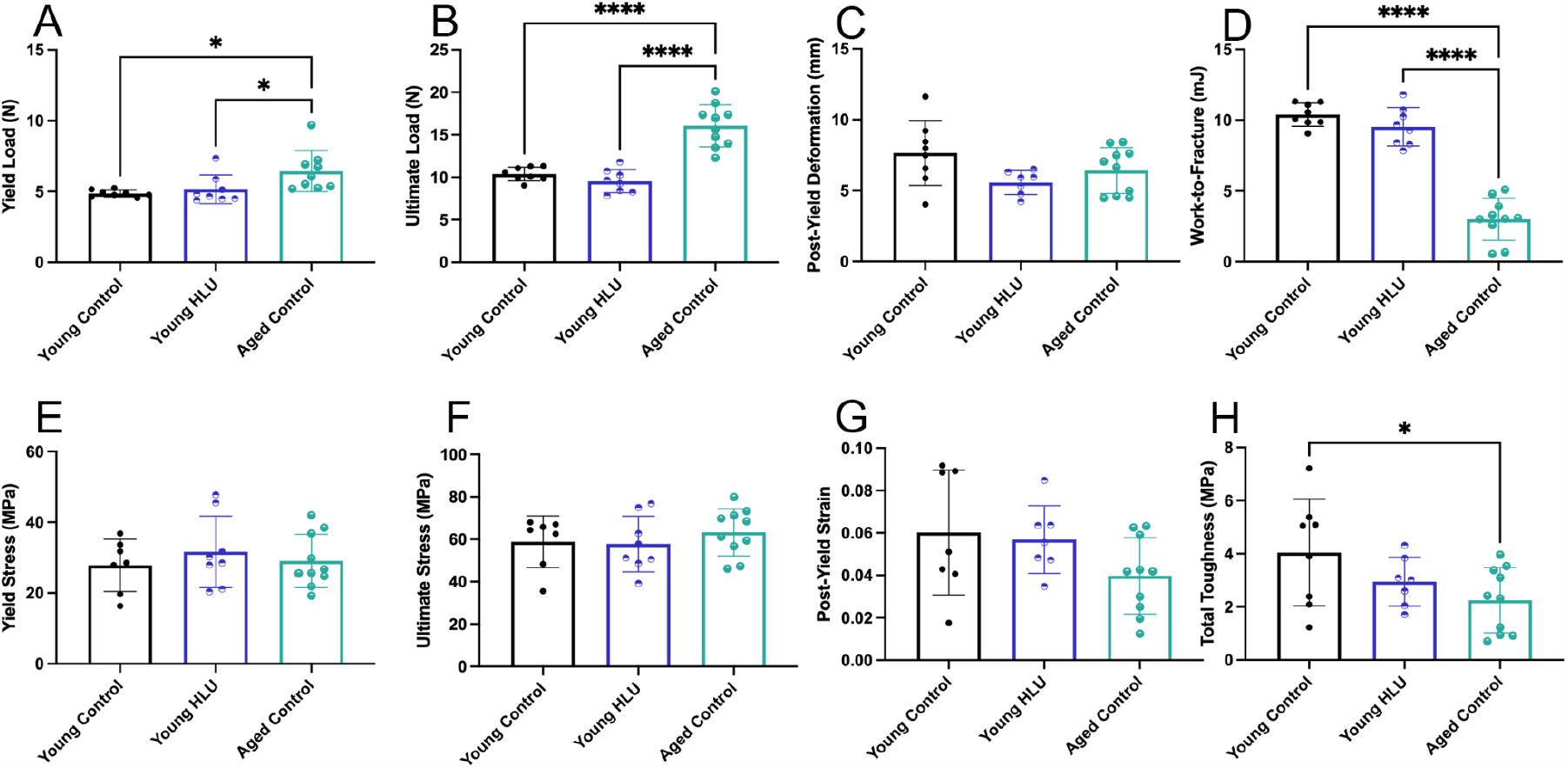
Three-point bending of right femurs in male and female mice. Yield load (A), ultimate load (B), post-yield deformation (C), work-to-fracture (D), yield stress (E), ultimate stress (F), post-yield strain (G) and total toughness (H). * p < 0.05, ** p < 0.01, *** p < 0.01, **** p < 0.001, n = 8-11

### Hindlimb unloading induces changes in gene expression that are age-dependent

After performing analysis on bone microarchitecture and biomechanical properties, we were interested in examining the gene expression profiles of each group. RNAseq was performed from marrow-flushed tibias in 4-month-old, 6-month-old and 22-month-old mice. Differential gene expression was performed and there were different patterns of gene expression was induced after three weeks of HLU in 4mo and 6mo (Figure 5). Principal component analysis was able to better separate groups based on age (Figure 5E) rather than HLU in either 4mo or 6mo (Figure 5A,C). Hierarchical clustering was performed to better identify the genes involved. In 4mo HLU vs control comparison, there were three main sample groupings and four gene clusters (Figure 5B). In 6mo HLU vs control comparison, there were four main groups and two gene clusters (Figure 5D). In 22mo vs 6mo, there were three main groups and two main gene clusters (Figure 5F). Cluster 1 from 6mo HLU vs control and Cluster 2 in 4mo HLU vs control and 22mo vs 6mo were similar. They primarily related to muscle contraction, actin-myosin filament sliding, muscle filament sliding based on GO Biological Process 2021, and impaired skeletal muscle contractility, abnormal sarcomere morphology, abnormal muscle physiology based on MGI Mammalian Phenotype Level 4 2021. These genes were highly expressed in 22mo in the 22mo vs 6mo comparison but had no clear pattern in either 4mo HLU vs Control or 6mo HLU vs Control. Cluster 4 from 4mo HLU vs Control, Cluster 2 from young HLU vs control and Cluster 1 from 22mo vs 6mo were similar. They primarily related to neutrophil mediated immunity, and more interestingly, inflammatory response and cytokine-mediated signaling pathway based on GO Biological Process 2021. MGI Mammalian Phenotype Level 4 2021 ontologies relate to increased susceptibility to bacterial infection, abnormal neutrophil physiology and abnormal immune system physiology. These genes were highly expressed in 6mo HLU vs control, with no clear distinction between groups in 22mo vs 6mo or 4mo HLU vs control. These findings indicate that there are changes in gene expression with HLU that may be age-dependent.

**Figure 5.**
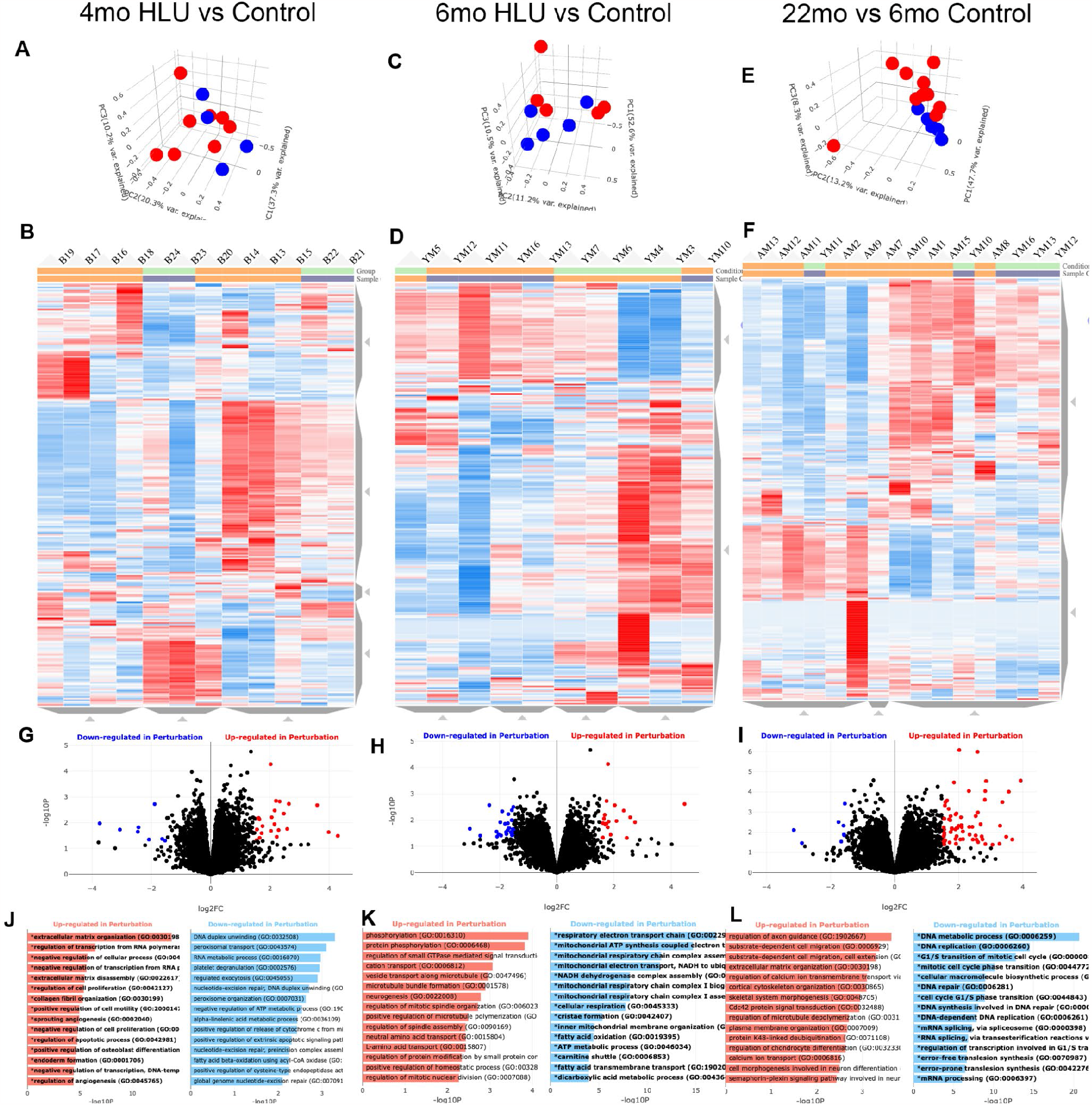
Processed RNAseq data from only male samples. Principal component analysis of RNAseq data comparing 4-month-old mice subject to HLU vs age-matched controls (A), 6- month-old mice subject to HLU vs age-matched controls (C), and 22-month-old mice compared to 6 month old control mice (E). Red = HLU or 22mo, Blue = Control or 6 mo, Clustergrammar of RNAseq data in each respective group. Green = HLU or 22mo, Orange = Control or 6mo (B, D,F). Volcano plot of each respective group (G, H, I). Red are significantly upregulated genes; Blue are significantly downregulated genes. logFC > 1.5, p < 0.05. Top upregulated and downregulated pathways based on gene ontology biological processes (2018 version). Bold indicates significantly enriched pathways. (J, K, L)

However, the gene expression changes are modest compared to changes that occur with aging.

Probing further into individual candidate genes, volcano plots are displayed in Figure 5G-I. Red dots indicate genes that are significantly upregulated in HLU or aged mice. Blue dots indicate genes that are significantly downregulated in HLU or aged mice. The threshold for significance was a logFC > |1.5| and a p-value < 0.05. There are more individual upregulated genes in 22mo vs 6mo compared to both 4mo HLU vs control and 6 mo HLU vs control, this is consistent with clustergrammar (Figure 5B.D.F) and PCA (Figure 5A,C,E) findings. Gene ontology was performed on the top upregulated genes (Figure 5J-L). In 4mo HLU vs control, notable ontologies include extracellular matrix organization, negative regulation of cellular process, and regulation of cell proliferation. In 6mo HLU vs control, notable ontologies include microtubule bundle formation, regulation of mitotic spindle organization, and regulation of mitotic cellular division. In 22mo vs 6mo control, notable ontologies include extracellular matrix organization, skeletal system morphogenesis, and plasma membrane organization. These results, interestingly, suggest that there is an increase in apoptosis and increased regulation of the cell cycle in response to HLU, these are characteristics also associated with age-related bone loss[2,26]. When examining the pathways related to downregulated genes, there are clues to internal cellular processes that become dysfunctional and lead to the downstream effects of each condition. In 4mo HLU vs control, notable ontologies include DNA duplex unwinding, peroxisomal transport and RNA metabolic process. In 6mo HLU vs control, notable ontologies include respiratory electron transport chain, mitochondrial ATP synthesis coupled electron transport, and mitochondrial respiratory chain complex assembly. In 22mo vs 6mo control, notable ontologies include DNA metabolic process, DNA replication, and G1/S transition of mitotic cell cycle. These ontologies suggest that HLU might cause dysfunctional mitochondria that can perhaps lead to cellular errors in transcription and translation, whereas in aging there is primarily a downregulation of cell cycle processes. These findings with aging are expected, given the importance of senescence with aging[27]. It is also interesting considering the different findings of HLU based on age, suggesting that there may be an age-dependency to the HLU response.

To further probe this age-related response to HLU, we included additional datasets that examined HLU vs. control mice. We assessed a female dataset from 3 month old C57BL/6 mice unloaded for 7 days[24]. The gene *Klb* was significantly upregulated in 2/3 unloaded age groups, and the gene *Sptlc3* was upregulated in 2/3 unloaded age groups. *Klb* codes for Klotho beta protein, predicted to be located in the plasma membrane or endoplasmic reticulum it is thought to be involved in fibroblast growth factor signaling, upstream of the MAPKKK cascade[28,29]. *Sptlc3* is a component involved in sphingolipid biosynthesis[30]. We also compared the total list of genes significantly upregulated in any HLU group with the list of genes significantly upregulated in aging. Most interesting was *Cyp2b10* (*Cyp2b6* in humans), a component of the cytochrome P450 system, which was upregulated in 6mo HLU and aging. There were no common significantly downregulated genes in any of the groups. These gene can be potential candidates for intervention of disuse-induced bone loss and to help identify the similarities between disuse-induced bone loss and age-related bone loss.

### Hindlimb unloading induced upregulation of genes involved in the senescence pathway in an age-dependent manner

Ingenuity pathway analysis was used to identify the leading canonical pathways and upstream regulators in each HLU age group (Figure 6). The top five canonical pathways based on z-scored identified by IPA were “Cardiac Hypertrophy Signaling”, “Senescence Pathway”, “Pulmonary Fibrosis Idiopathic Signaling Pathway”, “Oxytocin Signaling Pathway” and “S100 Family Signaling Pathway”. IPA identified “Senescence Pathway” as the second leading canonical pathway enriched in mice exposed to HLU. HLU induced activation of the senescence pathway in 3 month (p < 0.05, z = 3.464) and 4 month (p < 0.001, z = 0.6) old mice, and inhibited it in 6 month (p < 0.01, z = -3.153) old mice. 22mo vs 6mo was included as a positive control as we hypothesized (p < 0.0001, z = 3.157). When hierarchical clustering was performed, the canonical pathways expressed by 3mo HLU vs control and 22mo vs 6mo were most similar. 4mo HLU vs control was the next most similar to this sub- grouping, whereas 6mo HLU vs control was the most distant.

**Figure 6.**
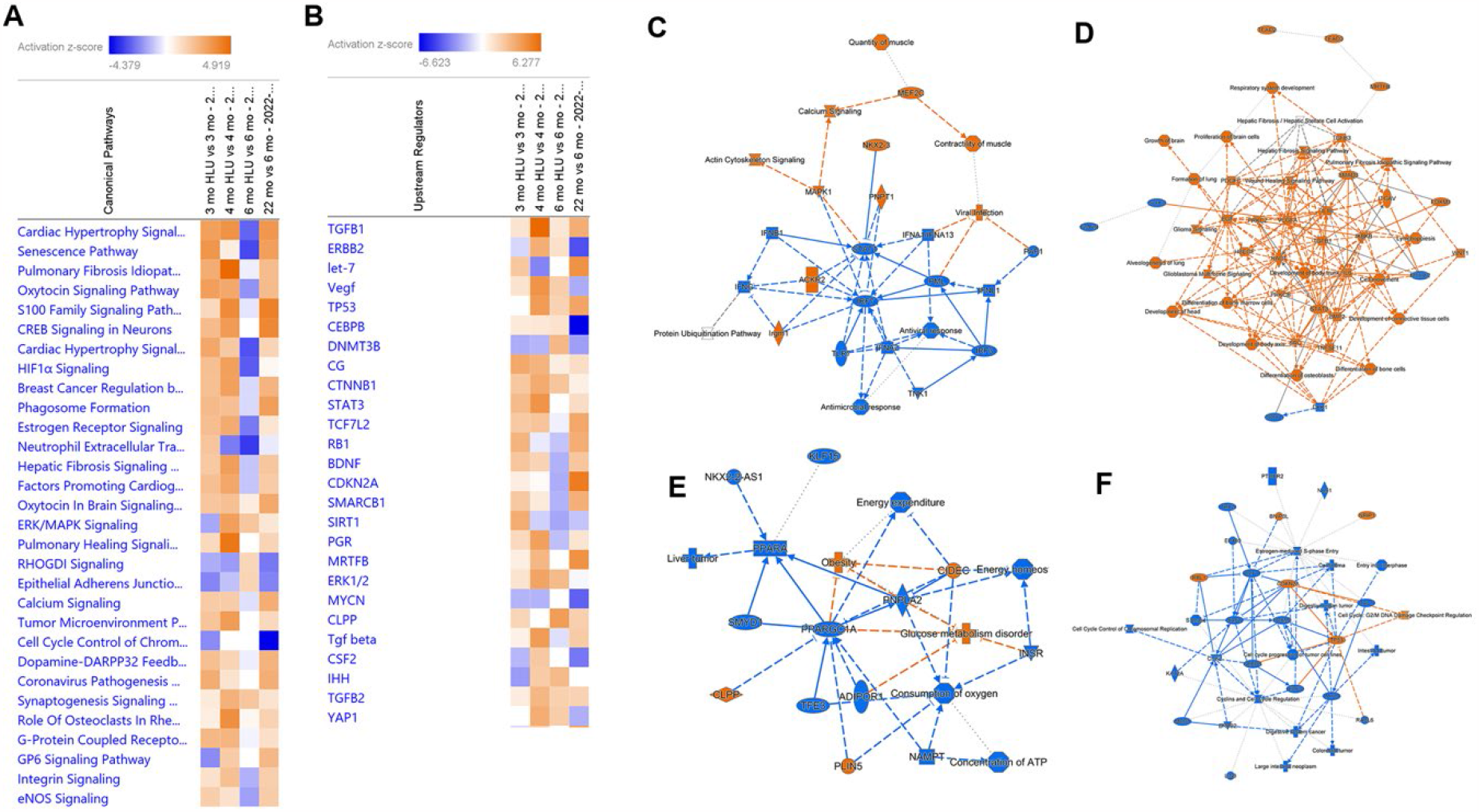
Ingenuity pathway analyses of RNAseq data from HLU vs control. Comparison analysis of main canonical pathways (A) and upstream regulators (B) in HLU vs control. (C) Summary of 3-month-old male mice in HLU vs control, (D) Summary of 4-month-old male mice in HLU vs control, (E) Summary of 6-month-old male mice in HLU vs control, (F) Summary of 22-month-old male mice vs 6-month-old male mice. Images are created using Qiagen’s IPA software.

The top upstream regulators in the “Genes, RNAs and Proteins” category predicted by IPA to be responsible for the pattern of gene expression were *Tgfb1, Erbb2, let-7, Vegf, Tp53 and Cebpb. Cebpb* encodes CCAAT/enhancer binding protein beta and is important in regulation of inflammatory gene products, including the senescence associated secretory phenotype[31,32]. IPA predicted *Cebpb* to be enriched in 3mo (p < 0.05, z = 0.897), 4mo (p < 0.0000001, z = 1.032), and 6mo (p < 0.05, z = 1.427) old mice exposed to HLU. *Erbb2* encodes erb-b2 receptor tyrosine kinase 2, a member of the epidermal growth factor family receptors and upregulation is associated with increased senescence[33]. IPA predicted *Erbb2* to be enriched in 3 month (p < 0.05, z = -1.057) and 4mo (p < 0.0000001, z = 3.402) exposed to HLU.

To determine whether changes in patterns of gene expression upon HLU were due to baseline changes in the senescence profile in age groups, IPA was performed to compare controls from each age (Figure S3). There was activation of the senescence pathway in 4mo (p < 0.05, z =3.677), 6mo (p < 0.01, z = 2.021) and 22mo (p < 0.001, z = 1.64) old mice relative to 3mo mice. There was similar activation of the senescence pathway in 22mo (p < 0.0005, z = 3.157) old mice relative to 6mo. Interestingly, there was decreased activation of the senescence pathway in 6mo (p < 0.00000001, z = -1.606) and 22mo (p < 0.0000001, z = -0.927) old mice relative to 4mo mice. Regarding upstream regulators, *Cebpb* (p < 0.00000001, z = -6.623) and *Erbb2* (p < 0.00000001, z = -4.548) were predicted to decrease in 22mo mice relative to 6mo mice, whereas *Cdkn2a* (p < 0.00000001, z = 5.487) was predicted to increase. In summary, in response to HLU in 3mo and 4mo mice the pattern of activation or inhibition of the most enriched canonical pathways and upstream regulators appeared to imitate the pattern in response to aging. However, this pattern was the opposite to the response to HLU in 6mo and 22mo mice. This suggests that there may be an age-dependent response to HLU.

Ingenuity pathway analysis was also used to generate summary graphics for each HLU vs control age group comparison and the aged vs young comparison (Figure 6C-F). Centrally located nodes have a higher number of connections than distally located nodes. In 3mo HLU vs control, there are genes related to a viral response. In 4mo HLU vs control, there are genes related to inflammatory cytokines *Il6, Il1b, Tnfsf11, Tgb1, Pdgfc* and their activated pathways *Stat3, Ikbkb, Smad3, Src*. There are also molecules involved in cellular proliferation and wound healing *Bmp2, Egf, Vegfa, Wnt1*. In 6mo HLU vs control, the central genes being influenced are *Ppargc1a* and *Pnpla2*, leading to dysfunction in energy homeostasis, consumption of oxygen and energy expenditure. The predicted disease states are glucose metabolism disorder and obesity. In 22mo mice vs 6mo mice, the central components of the interaction pathway relate to *Tp53, Cdkn2a, Rbl1* and Cell cycle: G2/M DNA damage checkpoint regulation. This variability in response to HLU in each age group suggests that there may be age-related differences after HLU in terms of the pathway leading to disuse-induced bone loss.

## DISCUSSION

Our hypothesis was based on the idea that age- and disuse-related bone loss shared a similar phenotype. In this study, we demonstrated that a disuse condition such as hindlimb unloading over a three-week period initiates several changes in bone microarchitecture and gene expression that is similar to aging.

This study was the first to directly compare the microarchitecture, biomechanical properties, and gene expression of young, unloaded, and aged mice to identify similarities. Trabecular bone compartments were more influenced by unloading than aging. This is interesting, considering that on visual inspection of microCT images there appears to be less trabeculae with aging, but this is not reflected on calculation. It is possible that software used to calculate trabecular bone volume was not able to accurately parse bone signals from background in aged mice due to the lack of trabeculae. Alternatively, the 7 μm voxel size that we used may not be sufficient resolution to observe the presence of thin trabeculae, at which point, nanoCT may be required to visualize changes. However, trabecular bone is important for transmitting forces throughout the anatomy of the bone, and so it would logically decrease in the context of disuse, but not aging[34,35].

Biomechanical testing results are interesting given the changes in bone microarchitecture. Most of the variability in bone strength is correlated with cortical bone parameters such as cortical fraction and cortical thickness[36,37]. Male mice demonstrated an increase in ultimate load and decrease in work to fracture with aging, but no change after disuse.

Gene expression results indicated that aging causes more transcriptome level changes than disuse. Our results indicate that aged mice already have significant defects in cell cycle processes, increased inflammation, and increased senescence. The cellular changes observed in disuse, particularly in regard to mitochondrial dysfunction and errors in the electron transport chain, are some signature features of senescence[38-40]. They can lead to increases in reactive oxygen species and activate senescence pathways[41,42].

Others have suggested that simulated microgravity *in vitro* induces senescence via mitochondrial dysfunction and oxidative stress in mesenchymal stem cells[43], skeletal muscle myoblasts[44], and neural crest-derived rat pheochromocytoma cells[45]. However, it should be noted that our model of disuse only occurred over three weeks. Senescence develops as a result of long-term chronic stress/insults, so it is possible that these observed changes are only early stages of dysfunction that can lead to senescence or apoptosis [6]. This might inform preventive strategies depending on timing of disuse. Perhaps prophylactic measures to prevent disuse-related bone loss should focus on mitigating mitochondrial dysfunction and oxidative stress, whereas chronic exposure may require a dual-treatment strategy against senescent cells and the SASP.

These data indicate that there is an age-dependent change in activation of the senescence pathway upon HLU in young adult mice (3-4 months), but not in slightly older mice (6 months). This work demonstrates that both aging and disuse upregulate the senescence pathway, but do not completely overlap. It is unclear why 4mo mice demonstrated the highest activation of the senescence pathway among the age groups in control mice. This time period coincides with peak bone mineral density and volume in C57BL/6 mice, after which there is a decline in cortical and trabecular bone architecture with increasing age[46]. It is possible that this arrest in gross bone growth coincides with a physiologic induction of senescence as observed during periods of embryonic development and wound healing[27,47]. These results will need to be experimentally validated to identify a mechanism that links mechanical stimuli to mitochondrial function.

It should be noted that only male mice were used in this study, except for the inclusion of GSE169292 (3month females) in IPA. Relationships in bone microarchitecture and biomechanical properties may be expressed differently in female mice. Female mice will need to be examined at these age groups to determine if there are sex-dependent effects when comparing disuse- and age-related bone loss[36,46].

Our results suggest that disuse may establish a senescent phenotype in young adult long bones. This has implications for young individuals who experience extended periods of disuse as would occur from lack of exercise, bedrest, limb immobilization and spinal cord injury. It also suggests that senolytics, drugs that selectively eliminate senescent cells, may be a future strategy to prevent disuse-induced bone loss in young adult long bones.

## Acknowledgments

This work was funded by NIH R01 AR068132, NASA 80NSSC18K1473, and CIHR DFSA 454761. We would like to thank the VCU Genomics Core for their assistance in performing RNAseq library preparation and sequencing.

