## Supplemental Files for "Hindlimb Unloading Induces Bone Microarchitectural and Transcriptomic Changes in Murine Long Bones in an Age-Dependent Manner"

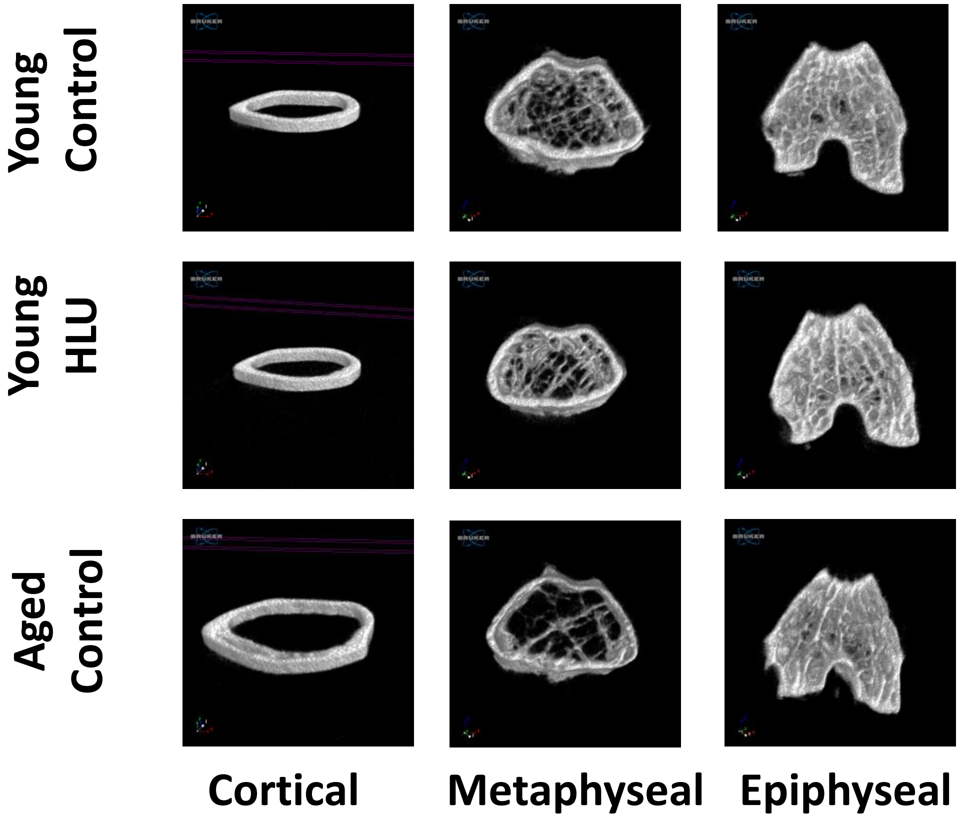


Figure S1. Three-dimensional reconstructions of left femurs after experimental intervention using microCT. Representative images from male mice.

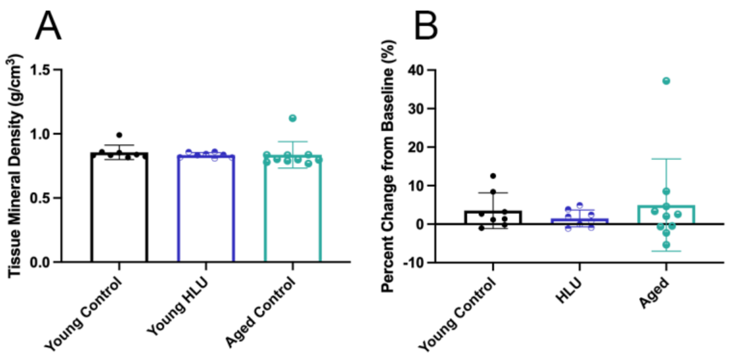


Figure S2. Bone mineral density changes in male and female expressed as absolute values after experimental intervention and percent change from baseline. * p < 0.05, ** p < 0.01, *** p < 0.01, **** p < 0.001, n = 3-15

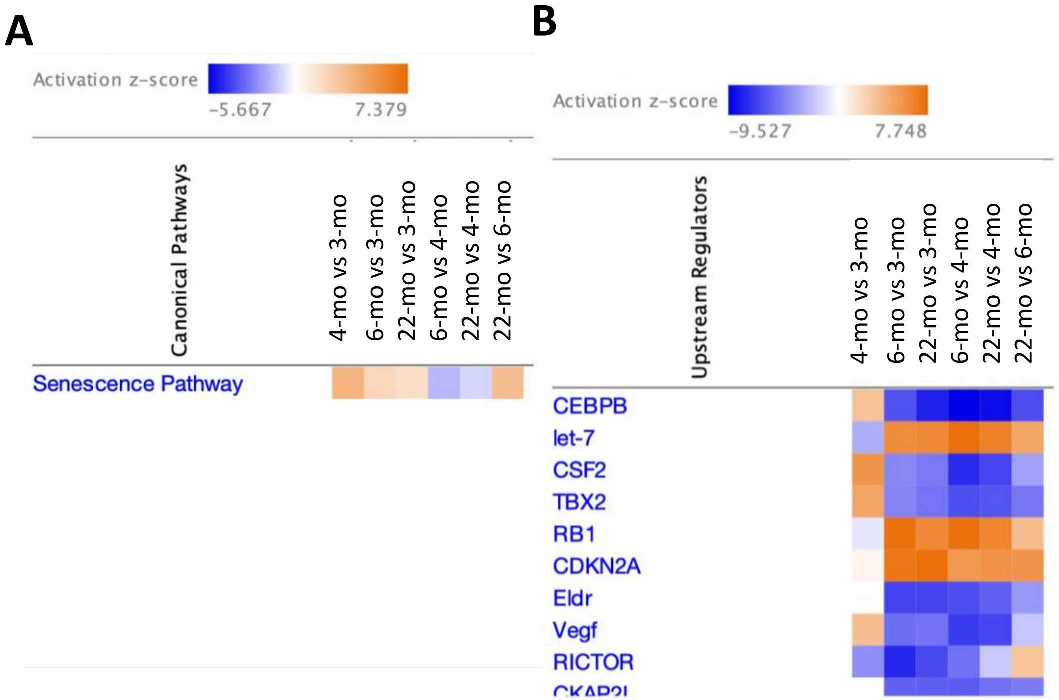


Figure S3. Ingenuity pathway analyses of RNAseq data from control groups. (A) Comparison analysis of main canonical pathways in control groups. (B) Comparison analysis of main upstream regulators in control groups.
